# Experimental susceptibility of North American raccoons (*Procyon lotor*) and striped skunks (*Mephitis mephitis*) to SARS-CoV-2

**DOI:** 10.1101/2021.03.06.434226

**Authors:** Raquel Francisco, Sonia M. Hernandez, Daniel G. Mead, Kayla G. Adcock, Sydney C. Burke, Nicole M. Nemeth, Michael J. Yabsley

## Abstract

Skunks and raccoons were intranasally inoculated or indirectly exposed to SARS-CoV-2. Both species are susceptible to infection; however, the lack of, and low quantity of infectious virus shed by raccoons and skunks, respectively, and lack of cage mate transmission in both species, suggest that neither species are competent SARS-CoV-2 reservoirs.

**Article Summary Line:** Experimental SARS-CoV-2 inoculation of North American raccoons and striped skunks showed susceptibility to infection, but transient, low-level shedding suggests that neither species is likely to be a competent natural reservoir.

## Introduction

SARS-CoV-2 continues to circulate on a global scale, the need to identify potential animal reservoirs, especially among wildlife, has become a priority, spurring surveillance and susceptibility trials of numerous species (*1,2*). Spillback infections from humans to animals has raised concerns about SARS-CoV-2 becoming endemic in native wildlife species (*3*).

The species belonging to Musteloidea (Mustelidae, Mephitidae, and Procyonidae) are ecologically and taxonomically relevant due to their high probability of becoming exposed to, infected with, and developing clinical disease to SARS-CoV-2 (*4*). For example, ferrets, close relatives to North American Musteloidea, are well-established animal models for coronaviruses (5) and are highly susceptible to SARS-CoV-2 (*6–8*).

Two Musteloidea, striped skunks (*Mephitis mephitis*, Mephitidae) and raccoons (*Procyon lotor*, Procyonidae), have established populations throughout much of North America, and are abundant, opportunistic, omnivorous generalists (9). Raccoons are also well established in regions of Europe and Asia (10). Both species have become habituated to food and shelter near human homes, resulting in frequent interactions with domestic animals, humans, and their waste. Skunks and raccoons are also notorious reservoirs of viruses that have substantial impacts on other wildlife and humans (i.e., rabies, canine distemper, protoparvovirus) (*11,12*). We therefore evaluated the susceptibility to infection, seroconversion, transmission potential between conspecifics, tissue tropism and pathology associated with SARS-CoV-2 in striped skunks and raccoons.

## The Study

We performed two independent susceptibility trails, first on raccoons and then skunks, using a similar experimental design. We directly inoculated 4 animals intranasally with 10^3^ (“low dose”) and 4 with 10^5^ (“high dose”) PFU of SARS-CoV-2 (direct inoculation, hereafter DI), strain SARS-CoV-2 USA-WA1/2020. This high dose has produced infections in ferrets and other species (*7,13*). The low dose was used to mimic the amount of virus to which these species may be naturally exposed (e.g., through consuming human garbage or potentially animal-to-animal) and has also resulted in infections and clinical disease in ferrets (*14*). DI animals were housed in pairs within ~1.5×1.5×2m wire mesh cages (Appendix Figure). We introduced a single, naïve conspecific to each pair at 48hrs after inoculation to test for direct contact (hereafter, DC) transmission. We housed 4 naïve animals of each species separately as controls and they were subjected to the same sampling routine as the experimental animals. Additional details are located in the Technical Appendix.

To document viral shedding, we collected nasal and rectal swabs daily from days post inoculation (DPI) 1-5, then on DPI 7, 9, 17 and DPI 7, 9, 15 for DI raccoons and skunks, respectively. DC raccoons and skunks were swabbed on DPI 1-5 and on DPI 7, 9, 11, 17 and DPI 6, 7, 9, 11, 15, respectively (Figure 1). Each sample was evaluated for infectious SARS-CoV-2 using virus isolation (VI) on Vero E6 cells and viral RNA (vRNA) using Real-Time Reverse Transcriptase PCR (rRT-PCR) that targeted both the N1 and N2 genes (*15*). All positive VI samples were confirmed with rRT-PCR. Viral titers were quantified using plaque assays. Shedding was detected from two high dose skunks on DPI 3, 4, and 5 and DPI 1 and 2, respectively. The highest shed titer was 3.3 log10/mL on DPI 4 (Appendix Table 1). Virus was not isolated from any raccoon or control animal samples. None of the DI, DC, or control animals of either species developed a fever, lost weight, changed behavior or displayed any signs of clinical infection.

**Figure 1.**
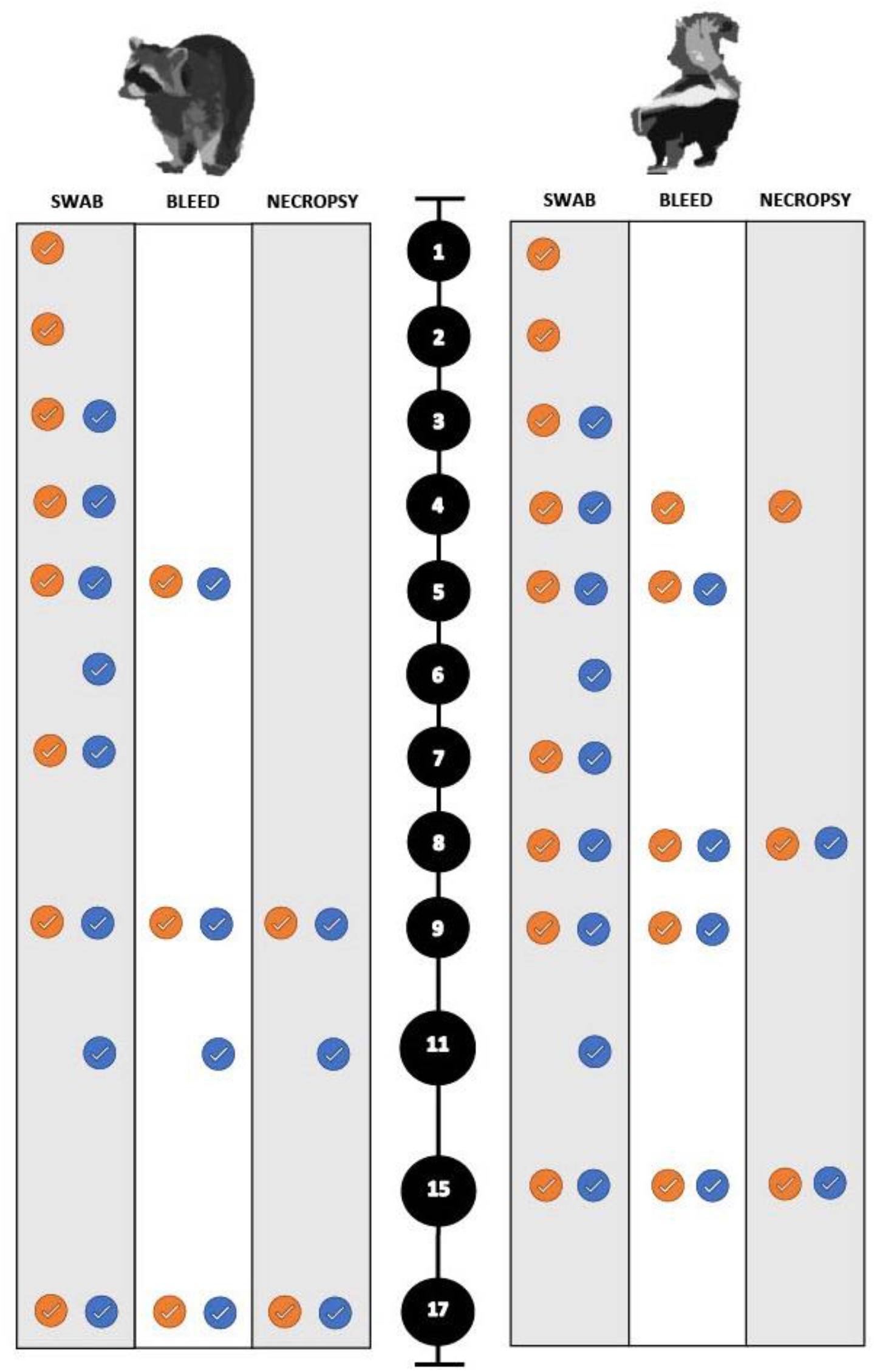
Timeline for the experimental SARS-CoV-2 infection trials of both raccoons and striped skunks. The black circles represent days post inoculation (DPI). The orange circles indicate directly inoculated animals. The blue circles indicate the direct contact animals.

All swabs and select tissue samples were evaluated for the presence of vRNA using rRT-PCR (cycle threshold (Ct) of ≤35). SARS-CoV-2 RNA was detected on the nasal swabs from 3 DI raccoons and 4 DI skunks; however, virus was only isolated from two of these skunks. One high dose skunk nasal conchae tissue sample was positive for vRNA. No virus was isolated from any of the tissues evaluated.

Blood samples were collected on DPI 5, 6, and 17 for raccoons, and DPI 5, 9, and 15 for skunks. Microtitration serum neutralization was used to detect seroconversion and determine endpoint titers. All DI animals of both species seroconverted, defined by a 4-fold increase in titers. The earliest seroconversion timepoint was DPI 9 for raccoons and DPI 8 for skunks, and the highest titers detected were 1:64 and 1:128, respectively (Appendix Table 2). No DC animals seroconverted, tested positive for the presence of vRNA, or shed virus.

Experimental animals were euthanized and necropsied at predetermined intervals to describe disease progression on histopathology and immunohistochemistry (IHC): raccoons at DPI 9, 11, 17, and skunks at DPI 4, 8, 15. Control individuals were euthanized and necropsied in tandem for comparison. Blood, nasal, rectal swab samples, and various fresh tissues were collected the day of necropsy. The gross necropsies for all animals were unremarkable. No microscopic lesions or SARS-CoV-2 specific immunohistochemical labeling were evident in tissues from raccoons.

The frontal and deep nasal conchae of 3/4 high dose, and 2/4 low dose skunks had mildly to moderately increased numbers of widely scattered lymphocytes and plasma cells in the superficial lamina propria vs. DC and control animals. At least one of four examined sections of lung of 3/4 high dose and 2/4 low dose skunks had mildly increased numbers of perivascular lymphocytes and plasma cells randomly scattered throughout the interstitium. There was no corresponding immunohistochemical labeling in these nor in any other tissues examined from skunks. All skunks had robust lymphoid tissue in lymph nodes, spleen, bronchus-associated lymphoid tissue (BALT; lungs), and gastrointestinal-associated lymphoid tissue (GALT; intestine).

## Conclusion

Our findings demonstrate that while striped skunks and raccoons are susceptible to SARS-CoV-2 infection, it is unlikely that either species is a competent reservoir for SARS-CoV-2 in a natural setting. No animals experienced clinical disease during this study. Raccoons exhibited no pathology, and skunks had mild evidence of subclinical recovering cellular response to viral infection in nasal conchae and lungs. The inability to isolate virus from raccoons, the lack of evidence of direct transmission between both species, and low amount of virus shed by skunks would likely impede the virus’s ability to establish in wild populations.

As with many susceptibility trials with wildlife, our animals were captive bred, healthy juveniles. Our finding may not readily translate into the susceptibility of wild populations due to factors such as senescence, immunocompetence (i.e., parasite burden, environmental conditions, gestation) and co-infections (e.g., distemper virus, parvovirus, etc.). Moreover, care should be taken to avoid transmission of SARS-CoV-2 to skunks and raccoons in a captive setting (e.g., zoological institutions, rehabilitation centers). Also, the rapid emergence of increasingly infectious SARS-CoV-2 strains in human populations presents new possibilities, such as increased transmissibility to marginally susceptible wildlife hosts.

While currently it seems unlikely for SARS-CoV-2 to circulate in raccoon and skunk populations and for virus to spillback into humans, other taxonomically related species, such as fishers, martins, and black-footed ferrets, especially those of conservation concern, have yet to be studied. Given that we did not document viral shedding, even in seroconverted raccoons, future wildlife surveillance studies should interpret antibody presence with caution. Continued global outbreaks of SARS-CoV-2 in farmed mink (*Neovison vison;(2)*) and spillback from humans to domestic and nondomestic animals highlight that additional susceptibility and transmission studies of Mustelidae are needed.

**Table 1.**
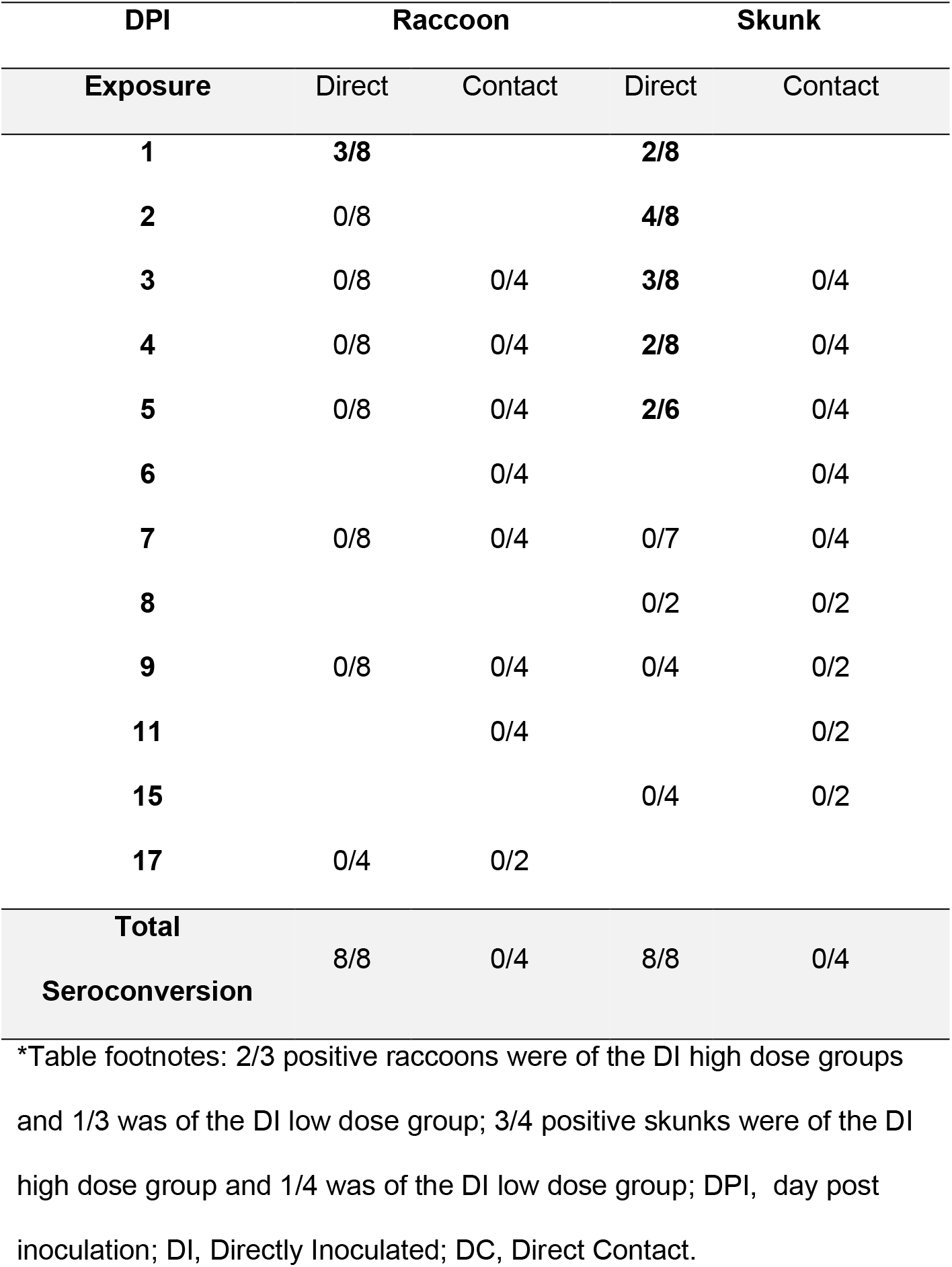
SARS-CoV-2 RNA detection in nasal swabs of intranasally inoculated and direct contact raccoons and skunks by real-time reverse transcriptase PCR (rRT-PCR).

## Supporting information

Technical Appendix

## Acknowledgments

We thank the technical assistance provided by the bioresources staff at the UGA Animal Health Research Center and histotechnologists at the Athens Veterinary Diagnostic Laboratory. We also thank Jeff Hogan, Department of Infectious Diseases, UGA for providing the SARS-CoV-2 isolate used in this study. Primary funding was provided by a National Science Foundation RAPID award (#2032044); student support was provided by an NSF EEID grant (#1518611). Additional funding was provided by the sponsorship of state fish and wildlife agencies to the Southeastern Cooperative Wildlife Disease Study (SCWDS). Support from the states to SCWDS was provided in part by the Federal Aid to Wildlife Restoration Act (50 Stat. 917).

## Author Bio

Raquel Francisco is a veterinarian and a Master of Science candidate within the Warnell School of Forestry and Natural Resources at the University of Georgia, USA. Her research interests include the inclusion of wildlife disease research to One Health strategies.

---

**Appendix Table 1.** SARS-CoV-2 virus isolated from the nasal swabs of intranasally inoculated striped skunks expressed in PFU (log10/mL).

| ID | Contact | DPI |  |  |  |  |
| --- | --- | --- | --- | --- | --- | --- |
| Skunks |  | 1 | 2 | 3 | 4 | 5 |
| H1-L | DI | 2 | 3.2 |  |  |  |
| H2-R | DI |  |  | 2.8 | 3.3 | 0 |

\*Table Footnotes: Both H1-L and H2-R were inoculated with a high dose (10<sup>5</sup>) of SARS-CoV-2. No animals were found to shed virus after DPI 5. PFU, plaque forming units; DPI, days post inoculation; DI, Directly Inoculated; DC, Direct Contact.

**Appendix Table 2.**
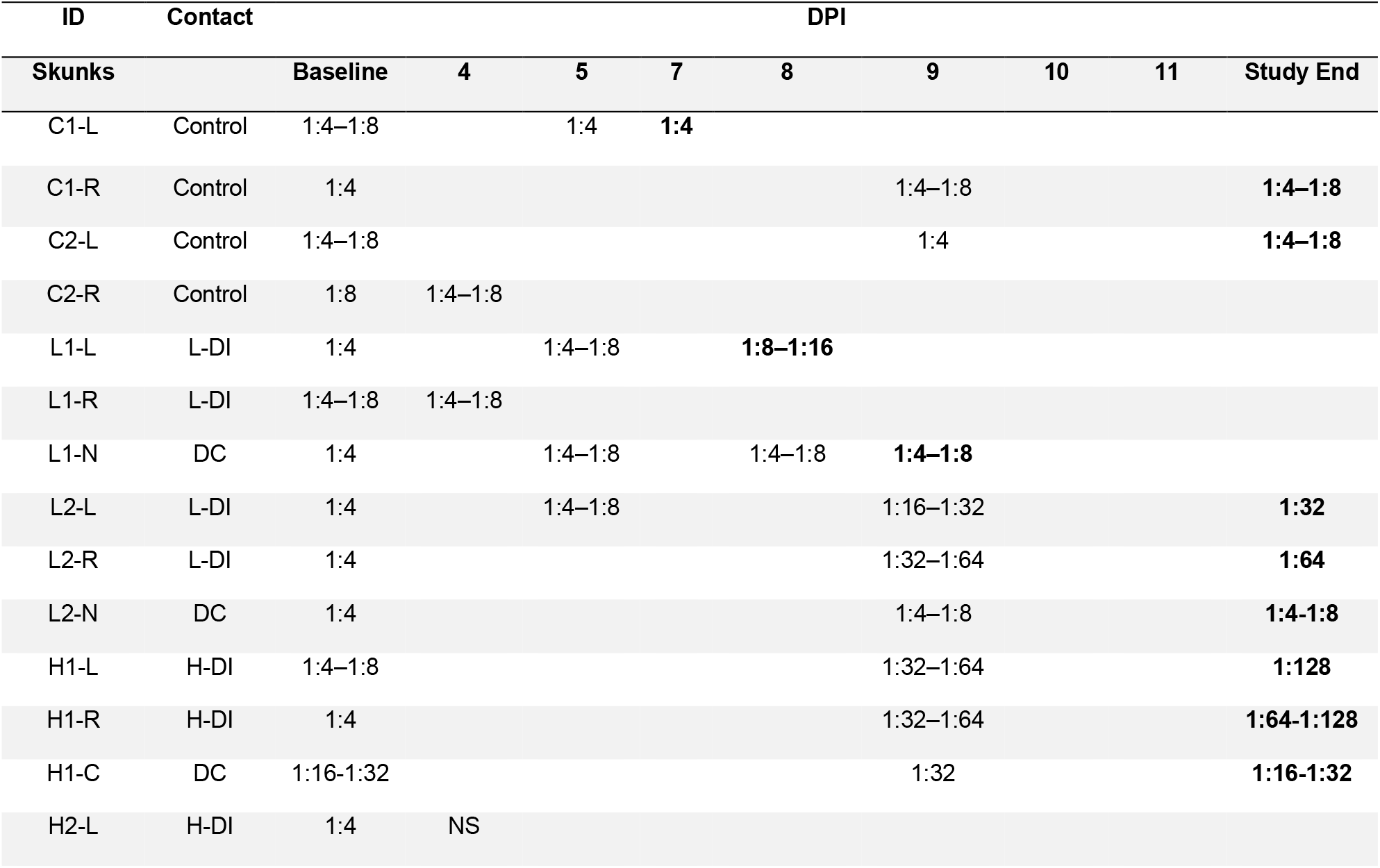
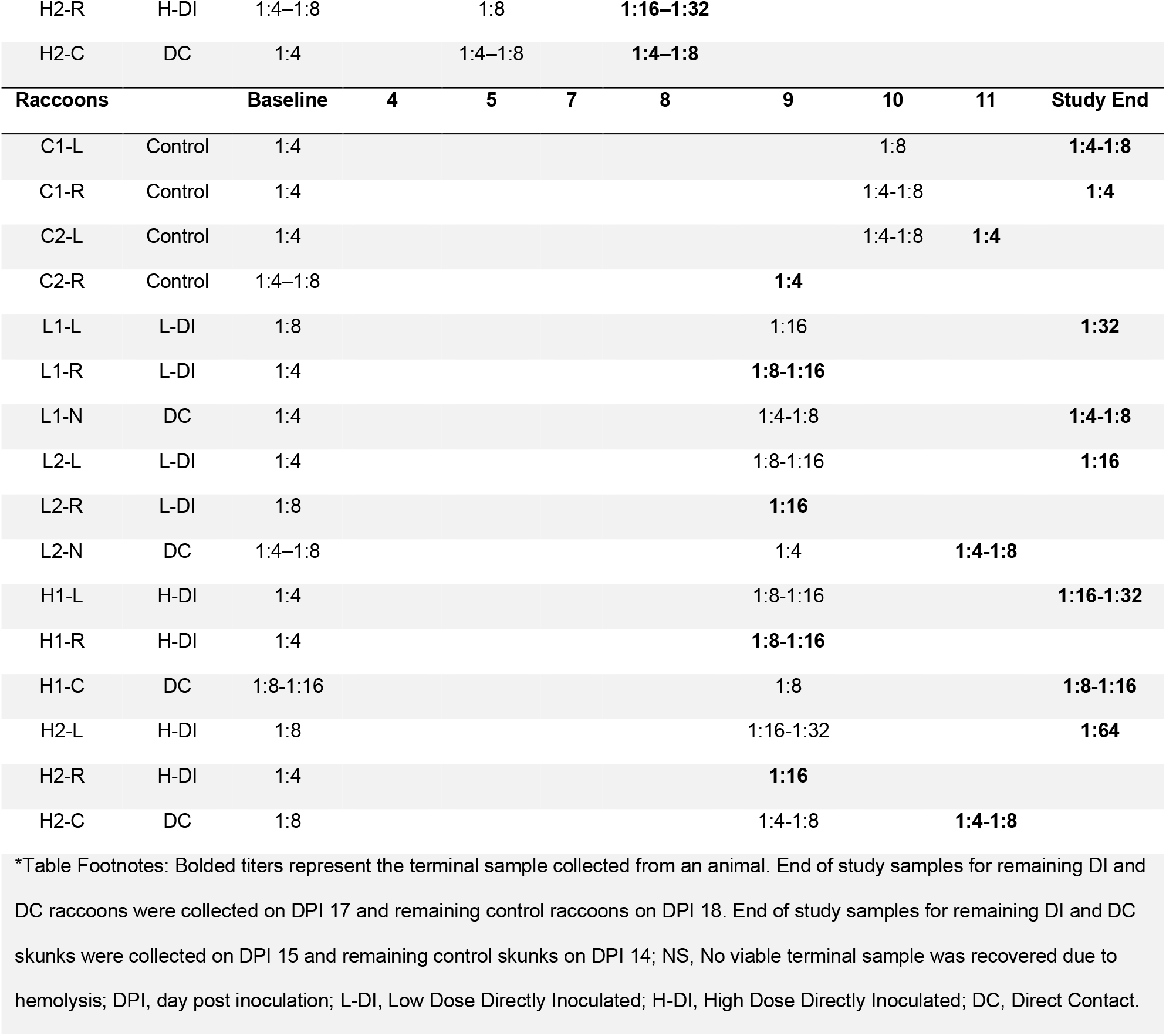
Serum neutralizing antibody development in striped skunks and raccoons intranasally inoculated with SARS-CoV-2, direct contact striped skunks and raccoons, and control striped skunks and raccoons.

**Appendix Figure.**
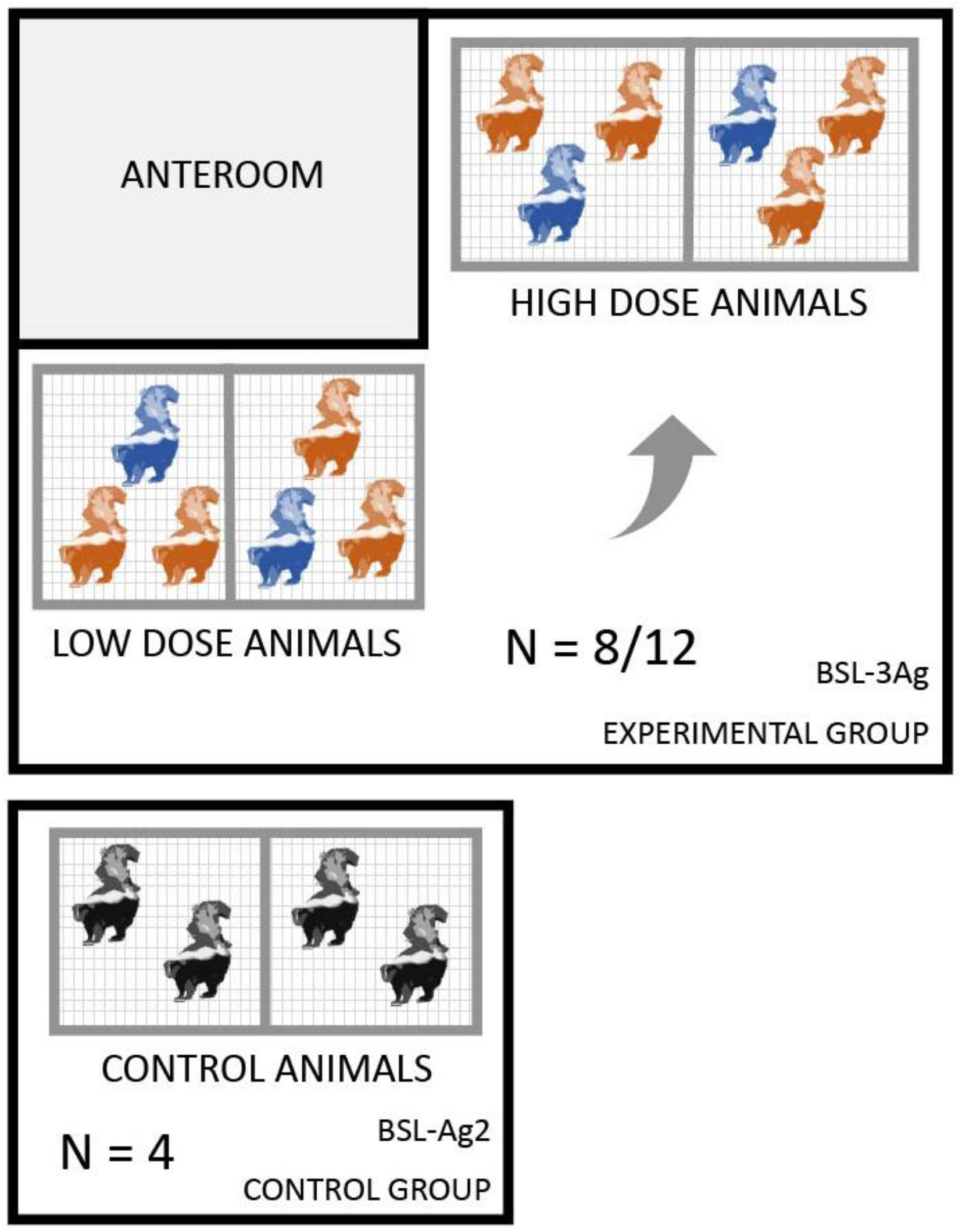
Agriculture Biosecurity Level 3 (BSL-3Ag) room layout for both raccoon and skunk infection trials. The orange skunks represent the directly inoculated animals and the blue skunks represent the direct contact animals. The room’s unidirectional airflow is represented by the arrow and did not recirculate.

