## Supplementary material for "Experimental susceptibility of North American raccoons (*Procyon lotor*) and striped skunks (*Mephitis mephitis*) to SARS-CoV-2": Technical Appendix

Technical Appendix and Other Materials

### Materials and Methods

Animals

Sixteen juvenile (~10 week old) male (n=8) and female (n=8), captive-bred raccoons and skunks were obtained from a commercial, captive breeding animal facility. All animal work occurred in the University of Georgia (UGA) Animal Health Research Center (AHRC) which is a high-security biocontainment facility. All of the work was conducted under Biosecurity Level 3 (BSL-3) procedures.

Animals were kept at ~21˚C and 50% humidity. Both species were fed daily with commercially available omnivore diet (Mazuri® Omnivore Diet, Purina Mills, LLC., MO, USA) and offered water ad libitum. The diet was supplemented by various fresh greens and protein items such as boiled eggs. Animals were identified by purposely trimmed patches of fur either on the left, right, or center of their rump.

Prior to inoculations, nasal swabs, rectal swabs, and blood samples were collected to ensure animals were not currently or previously infected with SARS-CoV-2. Samples were tested using virus serum neutralization test (VNT) and virus isolation (VI).

Ethics Statement: All procedures involving raccoons and skunks were reviewed and approved by the University of Georgia's IACUC committee (A2020 04-016) and Office of Biosafety (2020 0048).

Experimental Design

Experimental animals (n=12) were separated into 2 experimental dosing groups with equal sexes per group. Each dose group consisted of four animals housed in pairs in two adjacent stainless-steel cages (~1.5x1.5x2m). Both experimental groups were housed in the same BSL-3 Agriculture (BSL-3Ag) room but were separated by approximately 6m and the directional air flow in the room flowed from the low to the high dose group. The design of the BSL-3Ag facility does not allow for recirculated air, facilitating 13 to 15 air changes per hour, thus the likelihood of aerosol transmission between each group is negligible. Control animals (n=4) were housed in a separate BSL-3Ag room. The low dose and high dose animals were intranasally inoculated with $10^3$ PFU and $10^5$ PFU of SARS-CoV-2, respectively.

Virus and Inoculations

The SARS-CoV-2 isolate used was USA-WA1/2020, originally isolated from a middle-aged male in Washington, USA who traveled to Wuhan China in January 2020. Skunks and raccoons were inoculated with 5th passage virus. The virus was grown in vero-E6 cells (American Type Cell Culture (ATCC®) CRL-1586) which were maintained in minimal essential medium (MEM, 5L deionized water, 48g of Minimal Essential Media Eagle (Sigma-Aldrich, Co., MO, USA.), 11.11g bicarbonate) supplemented with 50mL/L of iron fortified calf serum (Sigma-Aldrich, Co.) and 20mL/L of Antibiotic Antimycotic Solution (10,000 units penicillin, 10mg streptomycin, 25µg amphotericin per mL). All cultures and microtitrations were incubated in a 5% $CO_2$ atmosphere and 37˚C.

For procedures, such as inoculation and venipuncture, raccoons and skunks were anesthetized with dexmedetomidine (0.04mg/kg) (Dexdomitor™, Orion Corporation, Espoo,

Finland) and butorphanol (0.2mg/kg) (Torbugesic™, Zoetis Manufacturing and Research, Girona, Spain) given intramuscularly (IM) then reversed using atipamezole (0.25mg/kg) (Revertindine™, Modern Veterinary Therapeutics, Germany) and naloxone (0.02mg/kg) (Wintac Limited, Bangalore, India) given IM (*1*).

The intranasal inoculations were performed on anesthetized animals using a 21-gauge catheter attached to a 1mL luer slip syringe (BD Syringe, Becton, Dickinson and Company, NJ, USA). Experimental animals that were intranasally inoculated with live virus (n=8) will be referred to as the Direct Inoculation (DI) groups. A single Direct Contact animal (DC) was introduced to each pair of DI animals 48 hours post-inoculation to evaluate direct transmission.

Sampling

Animal health status (i.e., mentation, attitude, physical appearance) was evaluated twice daily. Blood samples and weights were collected from raccoon experimental groups on DPI 5, 6, and 17 and control animals on DPI 6, 10, and 18. For skunks, blood samples and weights were collected from experimental groups on DPI 5, 9, and 15 and control animals on DPI 5, 9, and 14. A 2mL blood sample was collected from the jugular vein and added to plain 3mL sterile vacutainer blood collection tubes (Covidien™, MA, USA).

Animals were sedated for collection of nasal and rectal swabs using 30-45 mg/kg trazodone (Cadila Healthcare Ltd., Ahmedabad, India) suspended in either water or equal parts of ORA-Plus® Oral suspending vehicle and ORA-Sweet® (Perrigo, MN, USA). Raccoon nasal and rectal swabs were collected daily from DPI 1-5. Swabs were then collected on DPI 7, 9, and 17 for DI raccoons, DPI 7, 9, 11, and 17 for DC raccoons, and DPI 6, 8, 10, and 18 for control raccoons. Skunk nasal and rectal swabs were collected daily from DPI 1-5 as described above.

Swabs were then collected on DPI 7, 9, and 15 for DI skunks, DPI 6, 7, 9, 11, and 15 for DC skunks, and DPI 7, 9, and 14 for control skunks. To evaluate environmental contamination, swabs of food and water bowls were obtained each sampling period prior to any animals being handled. Swabs and blood samples were also collected for all animals when euthanized, regardless of group.

Nasal swabs were obtained by swabbing both sides of the nasal passage using a single sterile polyester swab (Puritan, ME, USA). Rectal swabs were obtained using sterile cotton swabs (Medline Industries, Inc., China). All swabs were placed in 1.5mL cryovials (SealRite®) with 1mL of Dulbecco's sterile phosphate-buffered saline (dPBS) (Sigma-Aldrich, Co.) for raccoons and 1mL of sterile virus isolation medium composed of MEM for skunks. The blood samples and swabs were kept cool until stored, and the whole blood was centrifuged within 1 hour for the collection of serum. All swabs and serum were then stored at -80˚C until processed.

Euthanasia

Animals were anesthetized, euthanized, and necropsied at predetermined intervals to maximize the chance of detecting histopathologic changes during the course of infection. All animals were sampled as described above after humane euthanasia. Animals were anesthetized with dexmedetomidine (0.04mg/kg) (Dexdomitor™), butorphanol (0.2mg/kg) (Torbugesic™), and ketamine (5mg/kg) (Zetamine™, OneVet, ID, USA), and euthanized with an intracardiac overdose of sodium pentobarbital (0.25mL/kg) (Euthanasia Solution, Med-Pharmex Inc., CA, USA).

On DPI 9, three raccoons (one low dose DI, one high dose DI, and a control) were euthanized and necropsied. On DPI 11 (i.e., 9 days post contact), two additional DC raccoons (one from each dose group) and a control raccoon were euthanized and necropsied. The remaining experimental and control raccoons were euthanized and necropsied on DPI 17 and 18, respectively.

On DPI 4, three DI skunks, (one low dose DI, one high dose DI, and a control) were euthanized and necropsied. On DPI 8 and 6, two DC skunks (one from each dose group), and two DI skunks (one from each dose group), were euthanized and necropsied. A control skunk was euthanized on DPI 7 for comparison. The remaining control and experimental skunks were euthanized and necropsied on DPI 14 and 15, respectively. Details on the sampling scheme of each species can be seen in Figure 1.

**Sample Analysis**

Virus isolation and Molecular Testing

Swab samples were placed in individual microcentrifuge tubes containing 1mL of viral media, vortexed, and then centrifuged at 10,000rpm for 10 minutes. Supernatant (100 µL) from each tube was inoculated into a separate well on a 12-well plate seeded with 3 to 4 day old Vero E6 cell culture monolayers. The plates were observed daily for cytopathic effect (CPE) for 10 days. If CPE was evident, the cell culture supernatant was collected and tested for the presence of SARS-CoV-2. Viral RNA (vRNA) was extracted from positive samples using the QIAamp Viral RNA Mini Kit (Qiagen Inc.), following the manufacturer's protocol. A validated real-time reverse transcriptase PCR (rRT-PCR) protocol was used for detection of SARS-CoV-2 (*2*,*3*). Reactions were conducted on a Step OnePlus Real-Time PCR System (Applied Biosystems, Inc.).

The same rRT-PCR protocol was used as described above to evaluate tissues and nasal, fecal, and environmental swab samples for the presence of SARS-CoV-2 viral RNA. A positive rRT-PCR result was defined as the detection of both the N1 and N2 genes. Both the N1 and N2 primer/probe had to have a cycle threshold (Ct) of ≤35 to be considered positive for the presence of SARS-CoV-2 RNA. Samples evaluated that resulted in a Ct of >35 for both probes were considered negative and samples with a Ct of ≤35 for one probe and a Ct of >35 for the other probe were also considered negative similar to Shriner et al(*4*). Viral stock with a titer of $10^5$ pfu/200ul was used as a positive control.

Skunk tissues (nasal conchae, tracheobronchial lymph node, tonsil, trachea (mid-length), lung (right middle lobe), heart, kidney, and jejunum) and select raccoon tissues (tracheobronchial lymph node, tonsil, lung (right middle lobe)) samples were homogenized with gentleMACS™ C Tubes (Miltenyi Biotec Inc., Bergisch Gladbach, Germany) using a gentleMACS™ Dissociator (Miltenyi Biotec Inc.). Tubes were then centrifuged at 3,220rpm for 10 minutes at 22˚C. Then, 100 µL of supernatant from each tube was inoculated into a separate well on a 12-well plate seeded with 3 to 4 day old Vero E6 cell culture monolayers. CPE was determined as discussed above. An additional 140 µL of supernatant from each tube was collected and tested for the presence of SARS-CoV-2 using the extraction protocol and rRT-PCR protocol listed above.

<u>Plaque Assays to Quantify Virus</u>

Cell culture supernatant (200 µL) from samples that were positive for SARS-CoV-2 via VI and rRT-PCR was diluted 10-fold (with the first well containing no dilution) for a series of 5 dilutions ($10^{-1}$, $10^{-2}$, $10^{-3}$, $10^{-4}$, $10^{-5}$), inoculated into a 6-well plate previously seeded with 4 day old Vero E6 cell culture monolayers, and incubated at 37°C and 5% $CO_2$ for 1 hour. Each well was then overlaid with 4mL of a gum tragacanth overlay solution (equal parts 2% gum tragacanth & 2XMEM, supplemented with 2mL of Fetal Bovine Serum (FBS), 5mL of Antibiotic Antimycotic Solution) and allowed to incubate as described above for 7 to 10 days. Once plaques were noted grossly, each cell culture was inactivated with 10% formalin and crystal violet solution and allowed to fix for 24 to 48 hours. Once cells were fixed, SARS-CoV-2 titers ($log_{10}$ PFU/mL) were evaluated in wells for which more than one plaque was present; no plaques were seen past $10^{-3}$ dilution on any sample. As previously determined, a ½ log is lost for each freeze thaw cycle, and all vials had been through 2 cycles, presumptively decreasing titers by 1 log (D.G. Mead, E.R. Lafortaine, unpub data).

Serology: Microtiter Serum Neutralization

SARS-CoV-2 neutralizing antibodies were detected and quantified using serum microneutralization. Serum samples were heat-inactivated at 56°C for 30 minutes. Then, samples were 2-fold serially diluted in duplicates from 1:4 to 1:256 and incubated at 37°C and 5% $CO_2$ with 100 $TCID_{50}$ of the same strain of virus used in the inoculum in 96-well plates for one hour. The wells were then overlaid with 150 µL of Vero E6 cells. The plates were incubated as described above and observed for CPE daily for 7 to 10 days, after which sample neutralization endpoint titers were determined.

Necropsy, Histology, and Immunohistochemistry

All inoculated and control animals were necropsied within 2 hours of euthanasia. Approximately 0.5 $cm^3$ samples of nasal conchae, tracheobronchial lymph node, tonsil, trachea (mid-length), lung (right middle lobe), heart, kidney, and jejunum were placed in cryovials and stored at -80°C for subsequent laboratory analyses.

Additional samples collected into 10% neutral buffered formalin for histopathologic evaluation included nasal sinus, trachea, lung (left cranial and caudal and right cranial and middle lobes), bronchus, lymph nodes (tracheobronchial, retropharyngeal, prescapular, and mesenteric), tonsil, tongue, esophagus, duodenum, jejunum, ileum, stomach, large intestine, liver (left lateral lobe), gall bladder, pancreas, spleen, heart, kidney, thymus, thyroid gland, adrenal gland, gonad, skeletal muscle (biceps), urinary bladder, bone marrow, cerebrum, cerebellum, brainstem, and eye. Once fixed, nasal sinus tissues were transferred to 12.5% neutral EDTA solution (250g EDTA disodium salt (J.T. Baker Inc. NJ, USA), 1750mL distilled water, and 25g sodium hydroxide) where they were allowed to decalcify for 14 to 21 days. Fixed tissues were routinely processed, embedded in paraffin wax, and 4µm thick sections were stained with hematoxylin and eosin (HE). Duplicate slides with deep nasal sinus, mid-trachea, left cranial lung lobe, bronchus, tracheobronchial and prescapular lymph nodes, tonsil, and jejunum for all raccoons inoculated with low and high SARS-CoV-2 doses also underwent immunohistochemistry (IHC) for SARS-CoV-2 antigen. These same tissues, in addition to frontal nasal sinus, left caudal lung lobe, right cranial lung lobe, retropharyngeal lymph node, kidney and heart also underwent IHC for all skunks inoculated with low and high doses, as well as the two high dose direct contact skunks.

IHC was performed on an automated stainer (IntelliPATH, Biocare Medical, Concord, CA). A rabbit polyclonal antibody for SARS-CoV-2 (ThermoFisher, PA141098) at a dilution of 1:100 for 60 minutes was used. Antigen retrieval on tissue sections was achieved using Citrate Solution 10X (BioGenex, Fremont, CA) at a 1:10 dilution 10 for 15 minutes at 110°C. A biotinylated goat anti-rabbit antibody at a 1:100 dilution (Vector Laboratories, Burlingame, CA) was utilized to detect the target, and immunoreaction was visualized using Warp Red Chromogen (Biocare Medical) for 10 minutes and counterstained with hematoxylin. A cell pellet with infected cells was used as a positive control.

The gross necropsies for all animals were unremarkable. No microscopic lesions or SARS-CoV-2 specific immunohistochemical labeling were evident in tissues from raccoons. The frontal and deep nasal conchae of high dose skunks: H1R, H2R, H2L and low dose skunks: L1R, L2L had mildly to moderately increased numbers of widely scattered lymphocytes and plasma cells in the superficial lamina propria vs. DC and control animals. At least one of four examined sections of lung of high dose skunks: H1R, H2R, H1L and low dose skunks: L2R, L2L had mildly increased numbers of perivascular lymphocytes and plasma cells randomly scattered throughout the interstitium. There was no corresponding immunohistochemical labeling in these nor in any other tissues examined from skunks. Incidentally, all skunks had moderate to severe, diffuse hepatic lipidosis and one low dose skunk, L1R, had focal, purulent rhinitis in the frontal nasal conchae.

All histology and immunohistochemistry were performed at the Athens Veterinary Diagnostic Laboratory at the University of Georgia and slides were read blindly by a board-certified veterinary pathologist.
